# Trade-offs in telemetry tag programming for deep-diving cetaceans: data longevity, resolution, and continuity

**DOI:** 10.1101/2022.05.28.493822

**Authors:** William R Cioffi, Nicola J Quick, Zachary T Swaim, Heather J Foley, Danielle M Waples, Daniel L Webster, Robin W Baird, Brandon L Southall, Douglas P Nowacek, Andrew J Read

## Abstract

**Background:** Animal-borne telemetry instruments (tags) have greatly advanced our understanding of species that are challenging to observe. Recently, non-recoverable instruments attached to cetaceans have increased in use, but these devices have limitations in data transmission bandwidth. We analyze trade-offs in the longevity, resolution, and continuity of data records from non-recoverable satellite-linked tags on deep-diving *Ziphius cavirostris* in the context of a behavioral response study of acute noise exposure. We present a strategic data collection programming scheme that balances resolution and continuity against longevity to address specific questions about the behavioral responses of animals to noise exposure in experimental contexts. We compare accuracy and precision between two programming regimes on a commercially available satellite-linked tag: (1) dive behavior summary defined by conductivity thresholds and (2) depth time-series at various temporal resolutions.

**Results:** We found that time-series data could faithfully replicate the more precisely defined dives from a dive summary record data stream to an acceptable error range for our application. We determined a 5-minute time-series data stream collected for 14 days balanced resolution with longevity, achieving complete or nearly complete diving records in 6 out of 8 deployments. We increased our data reception rate several fold by employing a boat based data capture system. Finally, a tag deployed in a group concurrently with a high resolution depth recorder showed high depth concordance.

**Conclusions:** We present the conceptual framework and iterative process for matching telemetry tag programming to research questions that we used and which should be applicable to a wide range of studies. Although designing new hardware for our specific questions was not feasible at the time, we were able to optimize the sampling regime of a commercially available instrument to meet the needs of our research questions and proposed analyses. Nevertheless, for other study species or designs, the complicated intersection between animal behavior and bandwidth of telemetry systems can often create a severe mismatch among research questions, data collection, and analysis tools. More flexible programming and purpose built instruments will increase the efficacy of these studies and increase the scientific yield relative to the inherently higher risk of invasive studies.

## Background

Satellite-linked bio-loggers (satellite tags) are an important component of the marine telemetry toolbox because they can uplink data to a satellite during deployments, and in some cases in near real time, rather than waiting for instrument recovery, which can be challenging for some species. Limited resources, uncertain detachment times, strong ocean currents, premature equipment failure, and battery life on radio beacons can all act to decrease the probability of device recovery. In fact, tags deployed ballistically on cetaceans often lack any flotation to keep size and mass low, and therefore there is no expectation of recovery. Nevertheless, bandwidth limitations are a serious drawback of many current marine satellite-linked tags, especially those that employ the Argos system, in at least two fundamental ways. First, the maximum bit rate such systems can support is constrained by hardware, orbiting characteristics, geographic coverage, and number of satellite receivers. Second, the behavior of the animal and device placement can be a limiting factor [1]. In cetaceans, this limitation is described by the frequency and duration of tag emergence into air, because successful uplinks occur only when the tag is out of water and a satellite is suitably nearby. Unfortunately, many of the marine species that are difficult to observe directly, and therefore of the greatest interest for bio-logging, are those that spend the least amount of time at the surface. Some of these species also dive to the greatest depths, which places additional stress on equipment.

A variety of methods have been employed to maximize the amount of data obtained from non-recoverable satellite-linked tags. Data can be compressed or summarized, duty cycled, and sampling rate and/or resolution can be lowered to better accommodate bandwidth restrictions [2]. Instruments can be designed to archive data, then release, and float to the surface where they transmit at a higher rate [3]. Finally, stations with ultra-high frequency (UHF) antennas and recievers affixed nearby to boats or on land can receive data, ameliorating the limitations of poor satellite coverage (for example, Argos Goniometer, [4]). Specific solutions depend on the behavior of the species of interest, the logistics of deploying receiving stations, as well as the data needed to address research questions [5].

We have been using several types of Argos-linked satellite telemetry tags to study the movements and behavior of *Ziphius cavirostris* (family: Ziphiidae) off Cape Hatteras, North Carolina, USA [6, 7]. Here we focus on SPLASH10 tags, which consist of a package of sensors including pressure, temperature, and conductivity, and an onboard computer and storage system that records, processes, and archives data. These tags are commonly attached to cetaceans in configurations that prevent recovery of the tag and the full archived record. In such cases, returned sensor data is entirely in the form of programmable data streams that are uplinked to the Argos satellite system, where they can be downloaded and decoded. A very commonly utilized data stream consists of dive summary records (termed *behavior* in the tag programming) for any dives which meet a predetermined threshold based on pressure, duration, and conductivity (user-definable within a certain range). Other gross metrics are available in data streams, for example depth histograms over a given time span (for example, daily). Finally, a true time-series of depths or temperatures can be recorded at one of five supported sampling periods and dynamically calculated data resolution.

In our applications, the surfacing behavior of the animals themselves creates a tremendous bandwidth bottleneck. The Argos system in use by these tags is limited to 32 bytes per data message and, although several messages can be sent per minute, typically only one message is sent during each surfacing of a whale. In addition, given the polar orbit of Argos satellites, at the latitude of our study site off the coast of Cape Hatteras, North Carolina (approx. 35–36^*°*^N), there is only approximately 9% temporal coverage. *Z. cavirostris* exhibit extremely long foraging dives (median: 59 minutes), shorter non-foraging dives (median: 19 minutes, and very short periods of ventilation during which they break the surface. The median duration of each ventilation period is only 2.2 minutes [7], during which time the animal will break the water’s surface multiple times.

Recently, we have successfully instrumented several dozen *Z. cavirostris* as part of the Atlantic behavioral response study, a strategic experimental design to quantify behavioral responses to naval sonar signals. The aim of this behavioral response study is to collect dive data before, during, and after known exposures to mid-frequency (3-4 kHz) active sonar (MFAS) signals using controlled exposure experiments either from operational Navy vessel-based sources or a simulated source [8, 9, 10]. These exposures are acute, up to 1 hour in duration, in select discrete time periods. Given this experimental design, we identified three key axes which represent trade-offs in satellite tag configuration for collecting dive data: overall data record length (longevity); temporal and spatial sampling scheme (resolution); and completeness or the number of gaps in the data record (continuity). The relative importance of these three trade-offs depends on the research question(s) being addressed, and an equal maximization function may not always be desirable. For example, prior to the start of the behavioral response study off Cape Hatteras, we programmed satellite tags to prioritize longevity and data resolution (Fig. 1) [7]. Specifically, we collected multiple data streams at relatively fine sampling rates, and duty-cycled data collection to increase battery life and overall transmission length, which necessarily introduced data gaps. In 2017, when the experimental phase of the study began, we wanted to determine whether mid-frequency active sonar disrupted deep foraging dives in known discrete exposure conditions, so we chose settings that prioritized longevity and continuity by collecting data only on foraging dives, at the cost of resolution. Thus, we employed a dive summary record only configuration, in which shallow dives were not recorded [2]. Later in 2018, we implemented a new programming scheme to address questions concerning potential behavioral responses over multiple behavioral states using the time-series data stream. In this scheme, we sacrificed overall longevity to produce a continuous record, centered around a known disturbance event of interest, over which behavioral state switching could be assessed.

**Figure 1.**
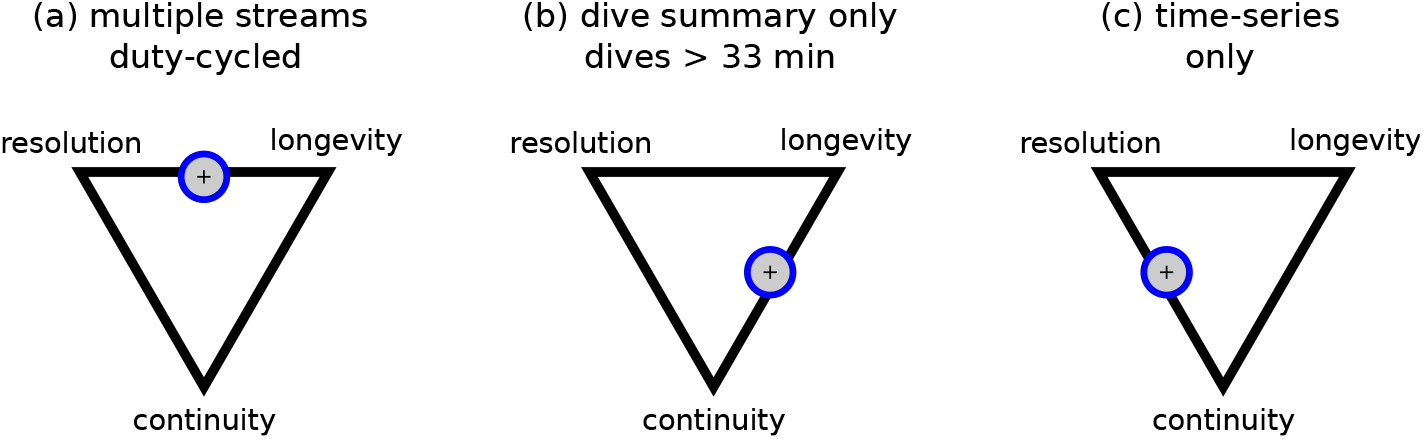
A conceptual framework of the trade-offs in bandwidth and battery limited bio-logging instrument programming. We highlight three axes: temporal and spatial resolution of the data, longevity of the data stream (and battery), and completeness of the data record. Position of the icon indicates roughly the priority balance for each of three setting regimes discussed here: (a) baseline settings with multiple data streams duty-cycled [7], (b) continuous dive summary records only [2], and (c) continuous time-series only.

Here we describe outcomes of increasing resolution and data continuity at the cost of longevity, given bandwidth limitations of Argos services, our field location, and *Z. cavirostris* behavior. We tested and deployed a programming scheme that employed only time-series data. In this paper, we report our programming optimization process within an experimental context, as well as a test of the efficacy of increasing bandwidth by employing an Argos Goniometer (Woods Hole Group, Woods Hole, Massachusetts, USA), a vessel-based UHF antenna and receiver system that intercepts radio transmissions from the tags. In addition, we provide practical insights for matching tag programming schema to specific research applications.

## Methods

### Tag deployment and programming overview

The data used in this analysis were a subset of the satellite tags deployed from 2014 through 2019 on *Z. cavirostris*. These instruments were satellite-linked depth-recording SPLASH10-292 tags with the extended depth range option (Wildlife Computers, Redmond, Washington, USA) in the LIMPET configuration [11] deployed using a DAN-INJECT JM 25 pneumatic projector (DanWild LLC, Austin, Texas, USA). Tags (*n* = 16) were attached with two 6.8-centimeter surgical grade titanium darts with backward-facing petals to the dorsal fin (*n* = 12), base of the dorsal fin (*n* = 2) or below the dorsal fin (*n* = 2).

We used two primary datasets in this analysis. The first dataset consists of 8 *baseline* tags (01–08) deployed in 2014-2016 programmed to collect a variety of data streams including both dive summary records (termed *behavior* in the tag programming) and time-series data at a 2.5-minute sampling period (Table 1). These tags were also configured to duty cycle for maximum longevity (see Supplementary Table 1, Additional File 1 for details). The second dataset consists of 8 *assessment* tags (09–16) deployed in 2018 with all optional data streams disabled, except for time-series depth measurements collected at a 5-minute sampling period. These samples are packaged in to discrete data messages containing information to decode 48 depth samples (4 hours worth). We programmed these tags to collect data only for the first 14 days of deployment, but to transmit data continuously (and therefore also generate position estimates) for the remaining life of the tag (Table 1). We chose 14 days because we estimated it would take approximately another 14 days to uplink a complete data record for a total of approximately 28 days. This total duration is similar both to the estimated battery life and our observed median instrument survival for SPLASH10-292 tags on this species [2, 7]. We deployed these tags as a specific test of the efficacy of this sampling scheme. We all use data from a single tag in the same configuration deployed in 2019 for the purposes of comparison with a higher resolution bio-logger deployed in the same group of animals (see below).

**Table 1.** Telemetry tag deployment summary. Purpose refers to the main use of the tags in the present analysis. DTAG pressure sensor data was decimated to 25 Hz before analysis.

| ID | Date | Longitude | Latitude | Tag type | Programming | Purpose |
| --- | --- | --- | --- | --- | --- | --- |
| Tag01 | 2014-05-15 | -74.78 | 35.55 | SPLASH10 | dive summary and time-series | baseline |
| Tag02 | 2014-09-16 | -74.71 | 35.66 | SPLASH10 | dive summary and time-series | baseline |
| Tag03 | 2015-06-14 | -74.74 | 35.60 | SPLASH10 | dive summary and time-series | baseline |
| Tag04 | 2015-06-14 | -74.69 | 35.63 | SPLASH10 | dive summary and time-series | baseline |
| Tag05 | 2015-10-15 | -74.77 | 35.61 | SPLASH10 | dive summary and time-series | baseline |
| Tag06 | 2015-10-21 | -74.75 | 35.62 | SPLASH10 | dive summary and time-series | baseline |
| Tag07 | 2016-05-27 | -74.74 | 35.59 | SPLASH10 | dive summary and time-series | baseline |
| Tag08 | 2016-08-21 | -74.69 | 35.61 | SPLASH10 | dive summary and time-series | baseline |
| Tag09 | 2018-05-24 | -74.78 | 35.69 | SPLASH10 | time-series only | assessment |
| Tag10 | 2018-08-05 | -74.78 | 35.73 | SPLASH10 | time-series only | assessment |
| Tag11 | 2018-08-05 | -74.78 | 35.72 | SPLASH10 | time-series only | assessment |
| Tag12 | 2018-08-05 | -74.75 | 35.55 | SPLASH10 | time-series only | assessment |
| Tag13 | 2018-08-06 | -74.78 | 35.48 | SPLASH10 | time-series only | assessment |
| Tag14 | 2018-08-07 | -74.78 | 35.57 | SPLASH10 | time-series only | assessment |
| Tag15 | 2018-08-07 | -74.78 | 35.56 | SPLASH10 | time-series only | assessment |
| Tag16 | 2018-08-07 | -74.75 | 35.59 | SPLASH10 | time-series only | assessment |
| Tag17 | 2019-07-30 | -74.73 | 35.54 | SPLASH10 | time-series only | DTAG comparison |
| Tag18 | 2019-08-06 | -74.75 | 35.58 | DTAG | pressure sampled at 250 Hz | DTAG comparison |

### Baseline comparison

We compared metrics derived from time-series and the dive summary record data streams on our 8 *baseline* tags where both of these data streams were at times running concurrently (Fig. 2, Table 1). The behavior summary data stream reports dive duration, maximum depth, and dive shape using a conductivity sensor to define the beginning and end of dives when a candidate dive passes a minimum duration (30 seconds) and depth (50 meters) threshold. Both maximum depth and a dive shape metric are calculated from a 1 Hz dive record with reported tolerance of 1 meter that is stored onboard the tag, but not transmitted to satellite. Three dive shapes are defined: square shaped dives were scored if greater than 50% of the dive was within 80% of the maximum depth, U-shaped dives were scored if between 20% and 50% of the dive was within 80% of the maximum depth, and V-shaped dives were scored if less than 20% of the dive was within 80% of the maximum depth [12]. Since the dive summary metrics are calculated from much higher resolution input data than the series data we considered the dive summary record data stream as truth and compared it to calculated metrics from the time-series data stream.

**Figure 2.**
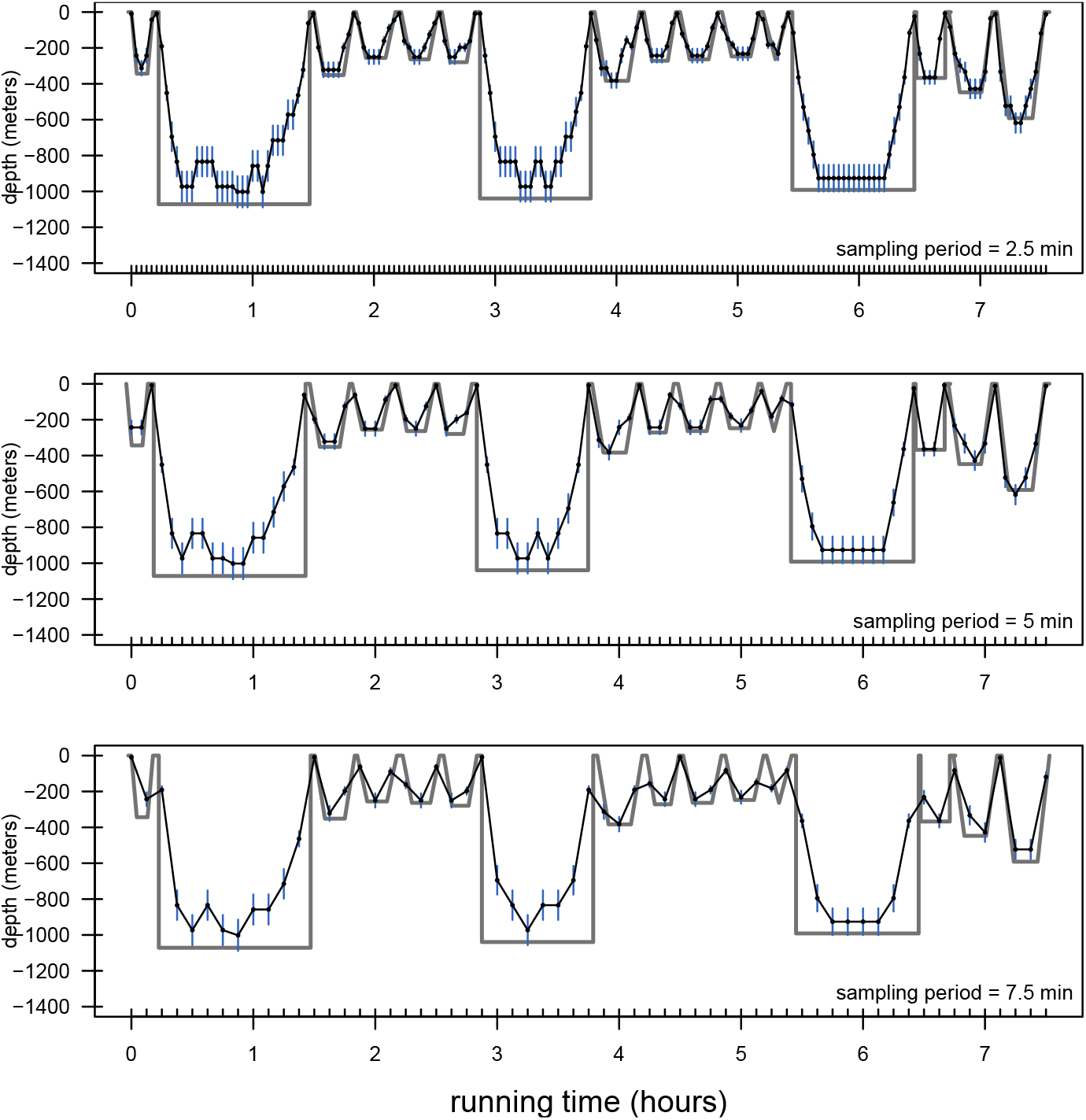
A representation of two types of dive data streams collected concurrently on a single tag deployed on a *Ziphius cavirostris*. In gray is a pseudo-dive profile based on a dive summary record data stream which provides a maximum depth, start time, and end time for each dive. Note this is not a true dive profile as only maximum depth and general shape of the dive are indicated (see methods for details). In black are time-series depths with reported error bands indicated by blue segments. Top panel shows the native resolution of time-series data for this deployment (period = 2.5 minute), while the middle and bottom panel show resampled time-series data (period = *{*5.0, = 7.5*}*).

We employed a simple algorithm to define dives and convert depth time-series data into a similar format as the behavior data, in which we used the first derivative of the time-series to determine points nearest a given surfacing with a depth filter of 50 meters to exclude local maxima. We visually inspected the resultant dives against the original time-series to correct any false negatives or positives. We interpolated surface times to calculate dive duration using a vertical velocity of 1.4 *m · s*^*−*1^ based on the findings of Tyack and coauthors [13], who found that ascent and descent rates varied little for *Z. cavirostris* within several hundred meters of the surface. Maximum depth was estimated from the largest value recorded between the start and end of the dive. We calculated dive shape using the same categories as the behavior summary data stream with the 2.5 minute depth samples as input. In addition, we resampled the time-series data from a 2.5-minute period to a 5.0- and 7.5-minute period to investigate the impact of sampling frequency (Fig. 2). We compared these converted time-series data sets to the dive summary record data on duration, depth, and shape that were recorded simultaneously on the tag.

We also compared depths from two different tag types deployed on different individuals in the same group in 2019, as animals in the same group at the surface are known to maintain high levels of synchrony [14, 15, 16, 17]. One of these tags was identical in type and programming to our 2018 time-series only satellite tags; the other was a shorter-term bio-logger attached by suction cups (DTAG) archiving pressure at 250 Hz, processed and decimated to 25 Hz [18].

### Additional data collected by UHF antenna

To aid in tracking and data collection we used an Argos Goniometer (Woods Hole Group Inc., Bourne, MA, USA; henceforth Goniometer) to localize tagged whales and receive data from their transmitters. We developed a visualization software in the form of an R package to assist in real time tracking of individual tags [19]. In addition, we used data messages downloaded by the Goniometer to supplement those data messages received only via satellite. We converted Goniometer-received hexadecimal data into a format that could be inputted into Wildlife Computer’s message decoding utilities using a custom R function [20]. Goniometer effort was approximated using the time difference between the first and last reception of a tag on the instrument per field day. Goniometers were affixed to 1 or 2 vessels per day which may have been engaged in a variety of activities including dedicated searching for previously tagged individuals.

### Assessment of time-series only configured tag deployments

Time-series only *assessment* tags (*n* = 8) were programmed with a 5.0-minute sampling period which was chosen to achieve the highest resolution possible with the most completeness, while giving up some longevity of dive data compared to our *baseline* tags. We measured the overall life of the tag from deployment to the final uplink, and the number of data messages successfully transmitted to satellite from the 14 days of time-series collection, and the number of consecutive messages without a data gap.

### Data validation

We checked for mechanical or software failures in our data streams. Status messages periodically report the pressure transducer reading at a presumed zero depth (when the conductivity sensor reads dry). We used these readings and manual inspections of the dive record to identify periods of excessive pressure transducer drift or failure. We defined unacceptably high pressure transducer drift as two or more consecutive absolute value zero depth readings of greater than 10 meters [21].

All analyses were carried out in the R programming language version 3.6.2 [22]. R packages **colorspace** 1.4-1, **ggplot2** 3.2.1, **reshape2** 1.4.3 and **R.matlab** 3.6.2 were used in visualizations [23, 24, 25, 26].

## Results

### Baseline comparison

We extracted a total of 645 dives from the 2.5-minute sampling period time-series data, compared to 598 and 457 for the 5.0- and 7.5-minute sampling periods respectively (*n* = 8 tags in all cases). Mean difference between time-series extracted dive duration and the dive behavior summary derived dive duration (error) increased as sampling period increased, although the maximum error was similar (Table 2). Most dive duration errors were within the theoretical maximum of twice the sampling period (Fig. 3). Time-series data tended to underestimate maximum depth, probably due to short forays to depths missed by the relatively coarse sampling. Mean depth error increased only very slightly with increasing sampling period, but the maximum depth error increased more substantially (Table 2). Correct assignment of dive shape also decreased with increasing sampling period from approximately 76% correct at 2.5 minutes to 65% and 63% at 5.0 and 7.5 minutes respectively. V-shaped dives were the most often miscategorized by the time-series data, but this type of dive was also the rarest. Square-shaped dives were the best identified at 76-79% correct for all sampling periods (Fig. 3).

**Table 2.** Mean absolute value differences among time-series data and concurrently collected dive summary records derived from the 8 *baseline* tags. Dive summary records include a dive duration calculated from submergence to emergence in air (as measured by a conductivity sensor) and maximum depth of each dive (as measured from an onboard pressure transducer sampled at 1 Hz). Time-series data were recorded at a 2.5-minute sampling period and resampled to 5.0- and 7.5-minute periods. Values in brackets indicate ranges.

|  | 2.5-min period | 5.0-min period | 7.5-min period |
| --- | --- | --- | --- |
| *n* dives extracted | 645 | 598 | 457 |
| duration (s) | 69 (0–1338) | 182 (0–1589) | 293 (0–1589) |
| depth (m), all dives | 30 (1–147) | 34 (1–147) | 43 (1–234) |
| depth (m), dives < 33 min | 18 (1–95) | 21 (1–101) | 27 (1–152) |
| depth (m), dives ≥ 33 min | 81 (2–147) | 82 (16–147) | 83 (16–234) |

**Figure 3.**
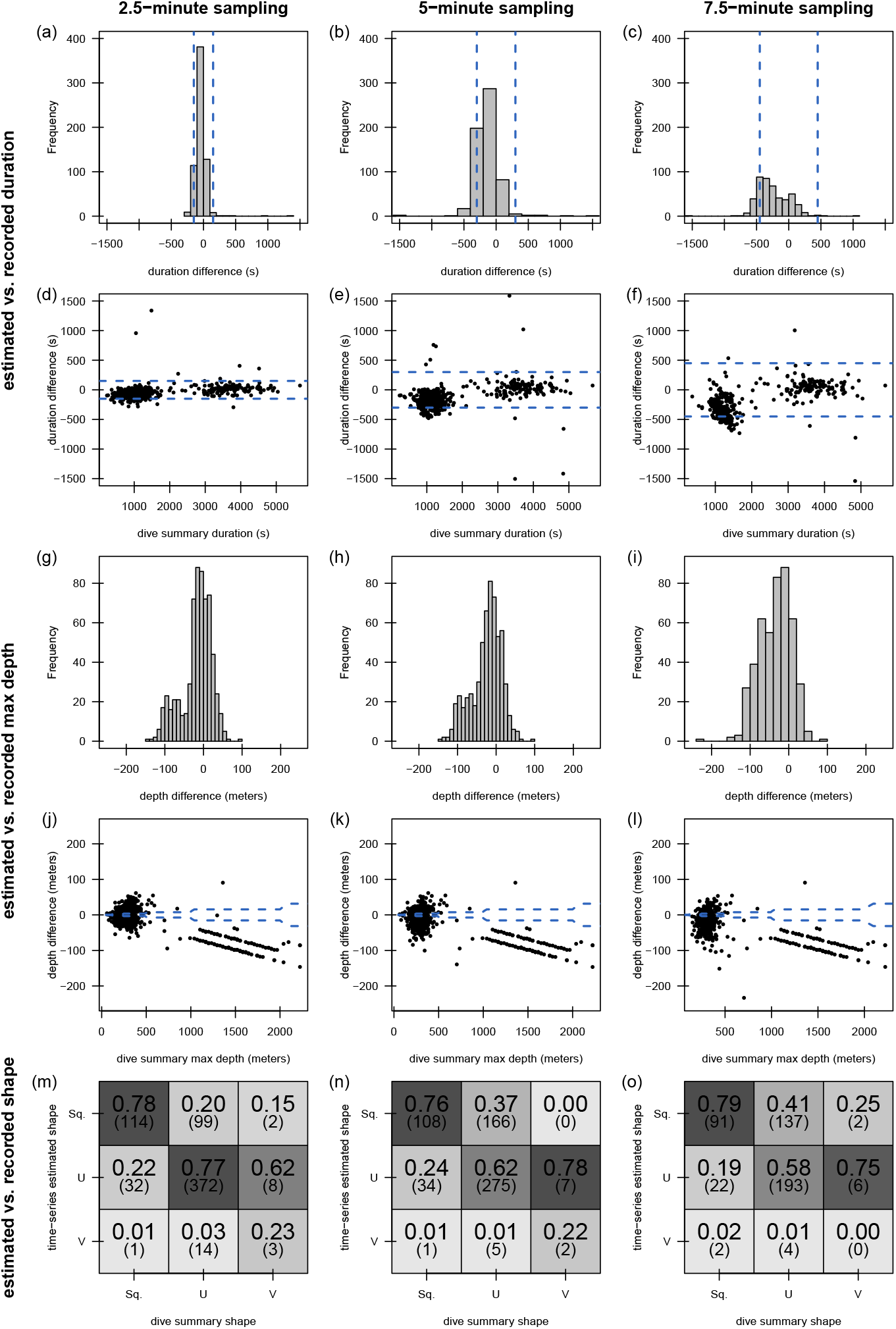
Dive metrics calculated from time-series depths recorded at a native sampling period of 2.5 minutes and resampled to 5.0 and 7.5 minutes compared to concurrently collected dive summary records derived from conductivity sensor detected dives. (a-f) show the distributions of the difference in dive duration between the two data collection methods for each sampling period. Blue broken lines indicate a theoretical error bound based on sampling period. (g-l) show the distributions of difference in dive depth between the two data collection methods for each sampling period. Blue broken lines indicate the recorded error bounds for the behavior summary data maximum depth. (m-o) show a confusion matrix of dive shapes calculated on board the tag (see methods for details) and transmitted in the dive summary records, and estimated post hoc from the time-series.

We also compared dive depths between the two whales tagged in the same group, which we expected to be highly synchronous. One of the pair was tagged with a highresolution DTAG and the other with a time-series programmed (5-minute period) SPLASH10 tag (Fig. 4). DTAG depth calibration error was 2.3 meters. Depths were highly correlated between the two instruments (*n* = 56 samples, *R*^2^ = 0.99) with a mean depth difference of 30 meters. Note that this difference includes both measurement error (from both tags), as well as any difference in the behavior of the two animals.

**Figure 4.**
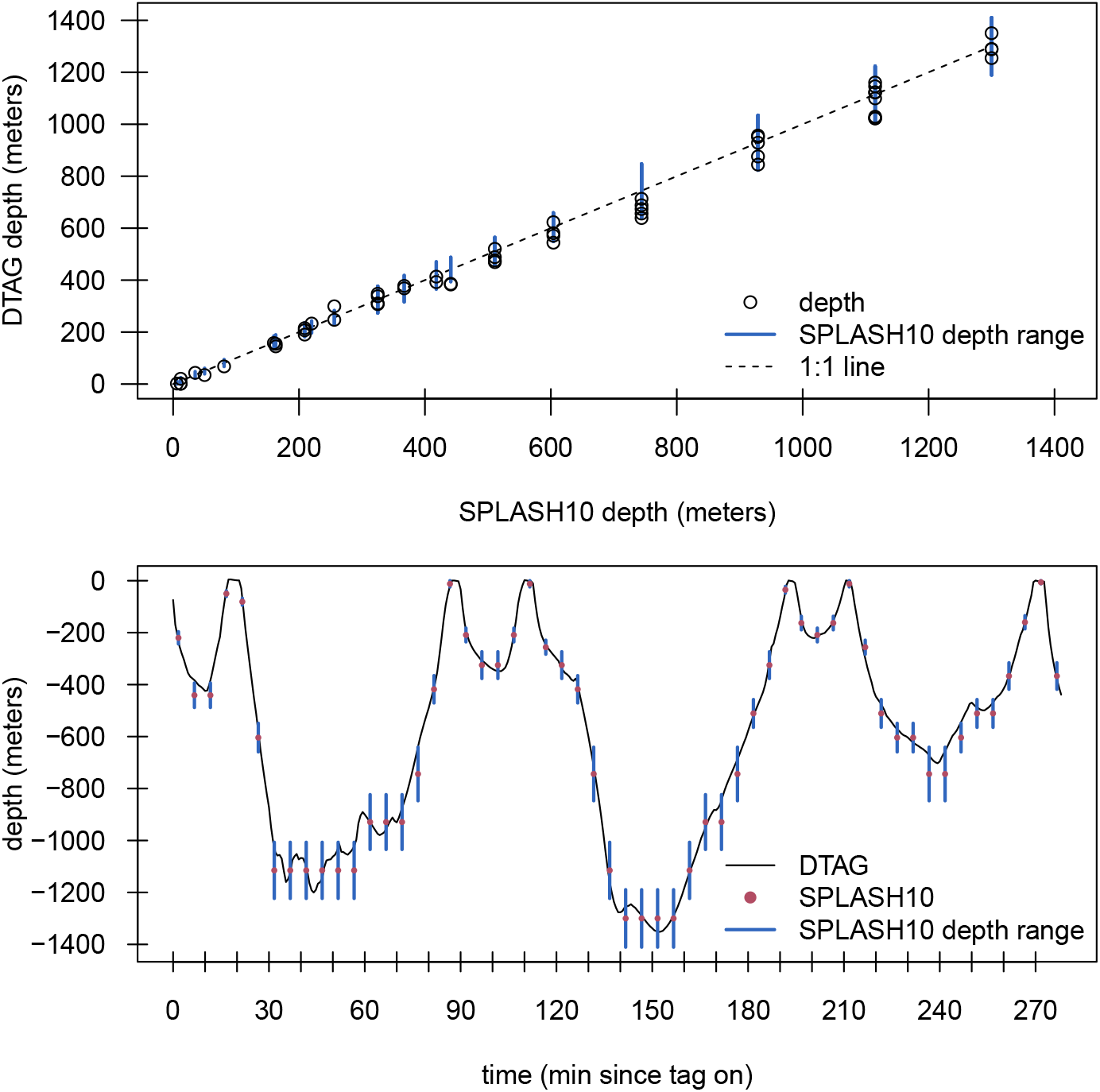
Comparison of dive depths between a satellite tag configured to collect time-series depth data and a DTAG deployed on different individuals in the same group. Top panel shows depth comparison with the broken line indicating 1:1 and gray segments indicating the reported error range for time-series reported depths. Bottom panels show the DTAG dive profile (Tag18) overlaid with 5.0-minute time-series depth sampling from the satellite tag (Tag17).

### Test deployments of time-series tags

We deployed eight time-series only tags to test the performance of this setting regime. We programmed these tags to collect 14 days of dive data, before transitioning to a transmit-only phase. One tag was deployed below the dorsal fin and never successfully transmitted any data (Tag10). A second tag suffered an unknown malfunction, apparently restarting at random intervals, which diminished the amount of transmitted data (Tag14). Of the remaining 6 tags, 2 experienced significant pressure transducer drift, resulting in truncation of the reliable data. These tags were still analyzed for data completion and transmission statistics because these aspects of tag performance were unaffected by the pressure transducer malfunction.

From the 6 tags that uplinked data, there were 3 data gaps across 3 different tags. Two of these gaps were 8 hours (2 data messages), while the last was only 4 hours (1 data message). Most data messages were received by satellite or by both satellite and Goniometer. Goniometer effort (time between reception of first and last Goniometer message) totaled approximately 212 hours over 31 days for our first vessel and approximately 178 hours over 26 days for our second vessel. Seven dive data messages (across 3 tags) were only received via Goniometer in the field (Fig. 5). The Goniometer also received additional status messages which did not reach satellites, although there was a higher rate of corrupt messages received in the field via the Goniometer than via satellite (Fig. 6). Daily rate of successfully decoded Goniometer messages was approximately 5 to 25 times greater than from satellite, while the corrupt rate ranged from 9 to 36 times greater.

**Figure 5.**
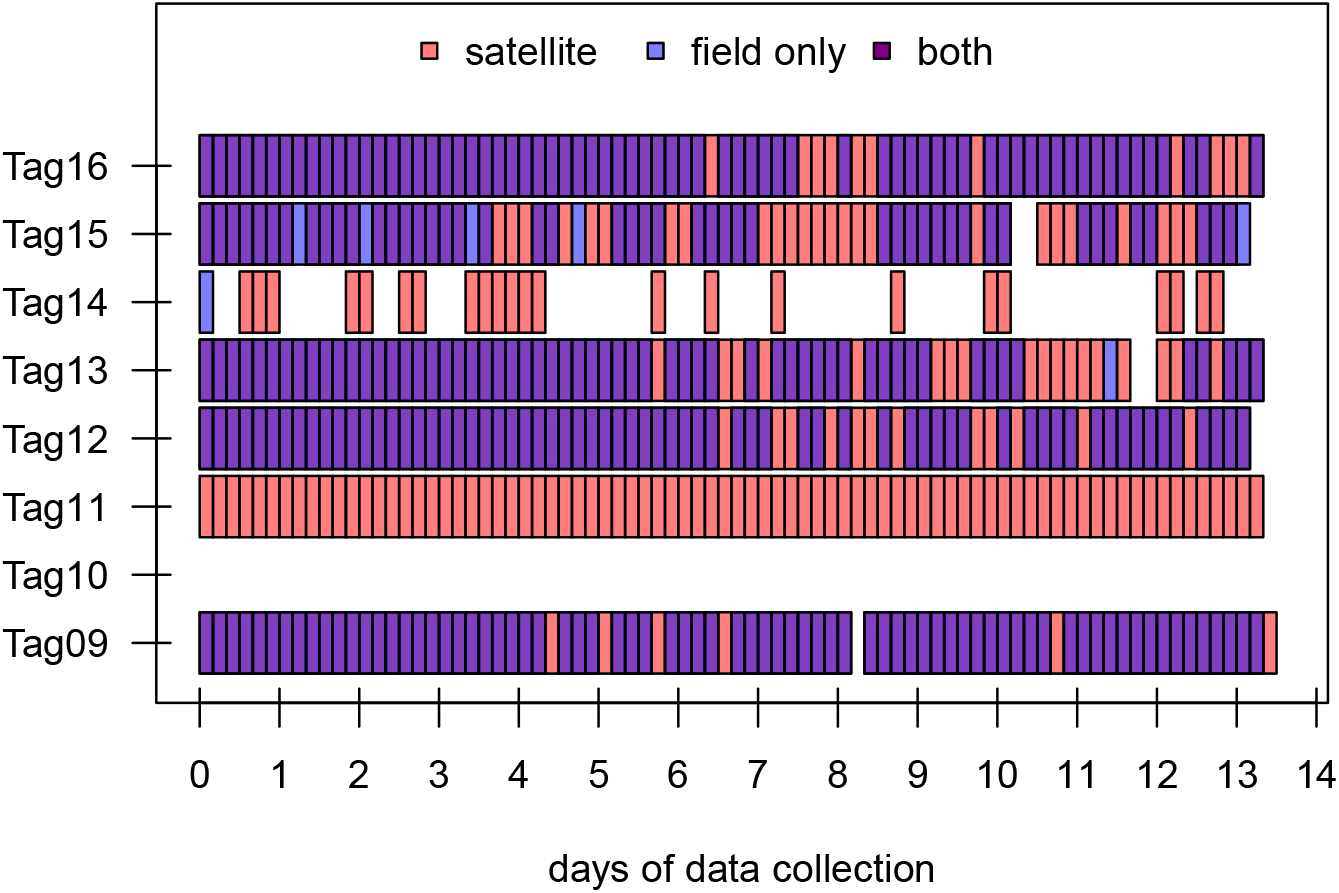
Time-series message reception for 8 test tags in the time-series only programming configuration. Each block represents 48 time-series data points (= 4 hours of data at our sampling period of 5 minutes). Colors denote if a message was received in the field only (via Argos Goniometer), from the Argos satellite system only or from both sources. Only successfully decoded messages are included in this plot. Tag10 never transmitted a successfully decoded time-series message. Note that total length of record varies as tags are programmed to record for 14 calendar days as opposed to exactly 336 hours.

**Figure 6.**
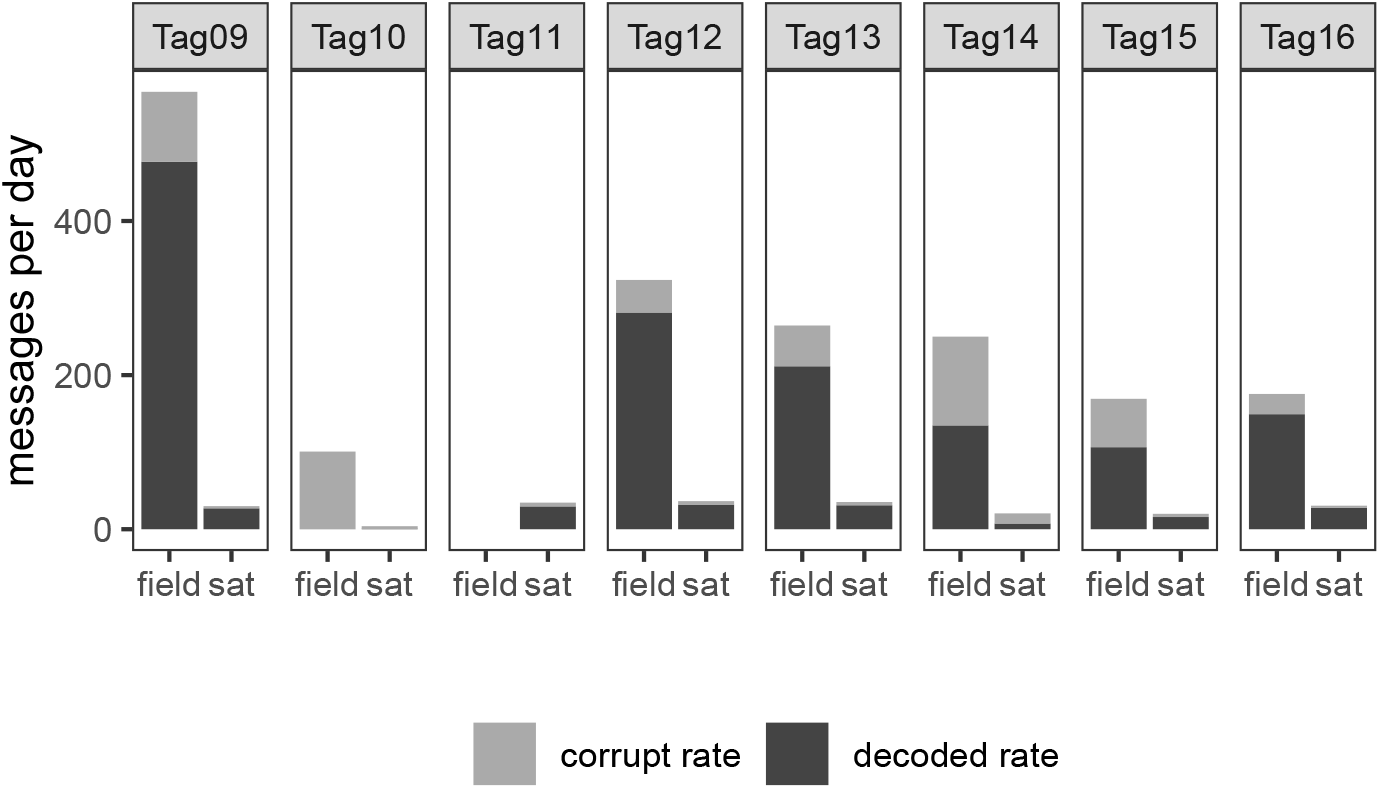
Comparison of tag message capture rate between the Argos satellite system (sat) and Argos Goniometer (field) for 8 test tags in a time-series programming configuration. The proportion of messages successfully decoded indicated by darker bars. Rates were calculated using the time from the first message received from a particular tag on a given day to the last message received that day. Tag11 was omitted from the field calculation, since it was only in reception range for a short period during deployment and not revisited.

## Discussion

Skin-piercing LIMPET configured telemetry tags are more invasive than many other possible data collection techniques and their deployment involves risk both to animal subjects and researchers [5]. Further, experimental behavioral response studies are by design invasive as a potentially harmful stimulus is introduced into the animals’ environment. Therefore every effort must be made to maximize the quantity and quality of data while minimizing impact and using the smallest possible number of animals. To those ends, this paper details the framework we used to match data collection to study questions, including when those questions evolved during the course of a single project.

### Sampling rate and data record length

We chose a 5-minute sampling period for time-series to maximize the temporal length of data, while reducing depth aliasing effects and minimizing data gaps due to limited uplink bandwidth. At the 5-minute sampling period, we estimate about twice as many data messages are generated per day than can be uplinked to satellite. We solved this problem by truncating data collection to 14 days, and then transitioning the tags to cease dive data collection but continue transmitting, enabling the tag to uplink the backlog of messages.

Relying on a post-data collection period to complete data transmissions also creates the risk that a tag could fail before the data uploading is complete. Our sampling length of 14 days was largely dictated by this concern. This total duration fell within the calculated battery life of the instrument, given our programming, and commensurate with average deployment life observed in our study area [7]. In addition, 14 days was sufficient to conduct experimental treatments including adequate baseline and post exposure periods given that experimental treatments were targeted for the middle of this period.

This sampling scheme was only viable for our experimental questions because the study species, *Z. cavirostris*, perform long deep dives, thus alleviating some of the depth and temporal resolution limitations of our tagging system. To answer similar questions about shallower, shorter or faster diving species or to answer questions about very small changes in depth, higher sampling rates and depth resolution would have been necessary. For some species and questions, an entirely different instrument may be necessary. Nevertheless, the approach we outline is applicable to those situations and would provide guidance on selecting the best data streams fit for a particular experiment or inquiry.

### Additional data collected by UHF antenna

To decrease the risks of not receiving data packages via satellites associated with this programming regime, we utilized a Goniometer to download additional data. Fortunately, by the end of our test deployments almost all messages that were obtained with the Goniometer had also been successfully received via satellite transmission, but the Goniometer provided additional security by collecting the most crucial messages (concurrent in time to the experimental exposures) before they were received by satellite and therefore guarding against potential future tag failure. Near realtime monitoring of received messages from experimental animals was possible in the field, which enabled strategic sampling to fill prioritized gaps and increased our ability to capture complete records. In the future, vessels with extended endurance (overnight capabilities) could greatly increase the potential data reception bandwidth, allowing for finer scale sampling or longer duration of sampling. These types of benefits have already been demonstrated in field sites with suitable land stations nearby [4].

### Iterative approach

There are clear trade-offs in any processes developed to optimize systems with known limitations, including the comparability of data among research efforts and the more limited time over which dive data are collected than would be possible at lower temporal resolution. Although we have focused on data resolution, longevity, and continuity, there are many other important factors to consider when deploying tags. These considerations include: weighing the risk of harm to the animal with the value of data collected [5, 27, 28]; the cost and time expenditure in deployment and analysis; how sample size is affected by programming regimes [29]; the appropriateness of data to biological questions [30]; species behavior; and the probability of success in achieving the experimental objectives during critical data collection periods.

One possible downside to tailoring tag programming regimes to each question or experiment is the complication of creating non-comparable datasets. For instance, if tags are deployed in the context of a long-term study, year-to-year comparisons may be of interest. For that reason, it is often more beneficial to collect data in a fashion such that it can be compared to historical samples, even as new questions and protocols are added to a project. In our case, the *baseline* data collection paradigm was not suitable to meet the specific experimental objectives from the Atlantic behavioral response study, given the short temporal (up to 1 hour) nature of experimental treatments [2].

### Added value of time-series data

Onboard data processing increases the efficiency of bio-logging devices immensely, especially when bandwidth is limited. For instance, when using the dive summary records to capture only long foraging dives, each data message comprises approximately 9 hours of *Z. cavirostris* behavior depending on the diving rate. In contrast, a time-series data stream set to a 5.0-minute sampling period only comprises 4 hours of data in a message and is dive rate independent. For species or applications where finer sampling is needed, this would be further reduced. In return, however, a true time-series even at relatively coarse resolution allows the calculation of activity budgets and summary statistics based on depth, spectral densities, custom shape parameters, and vertical velocity. These data are also well suited for more sophisticated continuous time behavioral modeling (for example, [31]). Again, our ability to recover this type of information from a relatively coarse diving time-series depends on the long deep dives of *Z. cavirostris* and sampling rate and depth resolution would need to be considered for other species and applications. Another benefit of the time-series in our case was that depth measurements were closely linked to a real-time clock, which was in contrast to the more temporally imprecise records in the dive summary record data stream. In the time-series, any concurrent tags sample almost simultaneously, allowing for direct comparison of the diving behavior of animals tagged within and between groups or, as we have shown here, even between different instrument types.

Another major consideration in our experiment was the depth resolution loss in the time-series only configuration. The lower depth resolution in the time-series was partially compensated by the fact that multiple depths were sampled during each dive, as opposed to a single depth in the dive summary record data stream (maximum depth) albeit at a finer resolution, but this could be an important consideration depending on the application.

### Species-specific behavior

The sampling resolution and depth accuracy to resolve, for example, individual dives is highly taxon-dependent, as is the degree to which the animal’s diving behavior creates bandwidth bottlenecks. *Z. cavirostris* create a significant bandwidth bottleneck by virtue of the small amount of time they spend at the surface, but the fact that their dives tend to be long and deep offsets this challenge by permitting the use of coarser sampling resolutions. Even the shorter dives of *Z. cavirostris* average 19 minutes [7], so a sampling period of 5 minutes does not typically cause aliasing, which could obscure dive events in the time-series record. Considering the sample period alone, it should be possible to detect any dive of 10 minutes or greater. Due to the limitations in depth accuracy, however, short and shallow dives may sometimes be unobserved. As the shorter dives of *Z. cavirostris* also tend to be relatively deep (*>*100 meters), this is not typically a problem for this species. In comparison, in the sympatric population of short-finned pilot whales (*Globicephala macrorhynchus*), the maximum recorded dive duration is 26 minutes and dives are typically shallower than for *Z. cavirostris* with a maximum recorded depth of 1360 meters [6, 32]. Therefore, a 5-minute sampling period would be insufficient to capture the same percentage of dives for this population. In fact for some applications this type of tag may not deliver suitable data at all for more shallowly diving species.

### Tag failure and limitations

We experienced multiple instrument failures during the deployments of the timeseries test tags. Three of the 8 tags suffered catastrophic failures rendering most of the return data unusable (a fourth tag was deployed too low on the animal to break the surface and transmit data messages). Such equipment failures are unavoidable in small-run electronics, especially when exposed to extreme conditions at or beyond their tolerances such as those deployed on deep-diving cetaceans, but failure rate must also be incorporated into the risk assessment of any programming scheme and, indeed, any tagging program. In this case, early failure could lead to dramatic reductions in completeness of the data record, so we took steps to mitigate this using the Goniometer.

Design limitations in our chosen instruments also impacted our data even when tags were functioning to specification. For example, the depth resolution of the time-series data in SPLASH10 tags is dynamically calculated from the maximum recorded depth (transmitted at some resolution itself) for a 1, 2, 4, or 8 hour data block (corresponding to the different sampling period options). Depths are split into 16 bins, which are narrower at shallow depths and wider near the maximum depth. This encoding is convenient since each depth point can be stored as just 4 bits, but can also cause complications in modeling as the resolution is constantly changing. In addition, the manufacturer declined to share the exact encoding algorithm, which further hampers efforts to produce consistent and reproducible analyses. There are similar drawbacks in the dive summary record data stream, such as a lack of precision in the real time code of dives (presumably to save bandwidth). The limitations mentioned here are device specific, but all instruments involve trade-offs in data collection choices, and these examples serve to highlight the general need to consider downstream data analysis before data collection especially in high risk projects and/or invasive protocols.

### Future development

There is a clear need for more flexible and/or purpose-built bio-logging instruments to answer many of the current and pressing questions in large marine vertebrate research, especially within the context of experimental behavioral response studies. In addition, there are specific requirements for instruments that are pushed to extreme environments, such as the significant pressure ranges visited by beaked whales. Hardware development is very expensive and, therefore, not always feasible, although in the case of deep-diving cetaceans at conservation risk, such research would be advantageous.

In our study, new hardware development was not possible, but we were able to optimize the sampling regimes of existing instruments available to us. Through this process it became clear more flexible and transparent hardware and software are needed. Additional control to set sampling rates and regimes could lead to more creative solutions in difficult bio-logging problems that would in turn enable data collection for a greater array of biological and applied conservation questions. Open source instruments could be a solution to creating accessible, flexible platforms for asking these questions consistently, transparently, and reproducibly and indeed these types of devices are on the rise (for example, [33]). This route will require strong partnerships between engineers and biologists (for example, [34]), and significant and ongoing commitment from funders.

## Conclusions

Lessons from our deployment and programming strategies should be generalizable to similar problems in other taxa and contribute to a growing literature on best practices in bio-telemetry. Our recommendations follow the logical thought process of any complex field experiment with specific objectives and constraints: start with the research questions, design analyses to address specific components, and optimize data collection for those analyses and questions. Testing data collection methods with pilot data or real deployments provides added value and allows for protocol refinement. Extensive testing, as presented here, is expensive and sometimes infeasible given the constraints of research budgets and the objectives of applied studies. Our funders allowed us to strategically and systematically evaluate tag settings to determine optimal solutions to best meet the specific research objectives of longterm studies of baseline behavior and behavioral responses of whales to sonar in our study site. The level of testing described here may not always be desirable and must be weighed with the potential impacts of an invasive instrumentation and the overall risk of a project. Computer simulations and bench tests are viable alternatives, but the intersection of animal behavior, weather, deployment location, and satellite coverage can be difficult to model or reproduce in the lab. A hybrid approach using simulation or modeling based on similar species and deployments can also increase the likelihood of success in field tests. Together these suggestions can serve to maximize scientific yield while seeking to minimize risk and impact to the study subjects.

## Abbreviations

UHF: Ultra-high frequency
MFAS: mid-frequency active sonar
LIMPET: Low Impact Minimally Percutaneous Electronic Transmitter

## Acknowledgements

We thank Joel Bell of Naval Facilities Engineering Command Atlantic for sustained project support and the latitude to experiment with methodology. Jessica Aschettino of HDR, Inc. provided assistance with tag deployments in 2018 as well as continued logistical support along with Daniel Engelhaupt of HDR, Inc. Jeanne Shearer prepared the DTAG data. We thank all members of the field team including Rafaella Lobo, Andrew Westgate, Jillian Wisse, Eleanor Heywood, Captain Reed Meredith of the F/V Kahuna, and Captain Jimmy Horning Jr. of the F/V Hog Wild. Greg Schorr and Dave Haas provided helpful input on tag programming and strategy. Matthew Rutishauser and Kenady Wilson of Wildlife Computers assisted during discussions on technical aspects of the tags.

## Authors’ contributions

AR, BS, DN, RB initially conceived the satellite tag deployment design (the behavioral response study) which generated these data and DLW, DMW, HF, NQ, WC, ZS provided additional input. DLW, DMW, HF, WC, ZS led the field work and data collection; AR, BS, DN, NQ, provided additional field assistance. WC conceived of the current study, analyzed the data, and wrote the first draft of the manuscript. All authors contributed to manuscript revision, read, and approved the submitted version.

## Availability of data and materials

All scripts and data used to produce these analyses and figures are available at https://github.com/williamcioffi/zc_series (doi: 10.5281/zenodo.6589596).

## Ethics approval

Satellite tags were deployed under National Marine Fisheries Service scientific research permit numbers 17086 and 20605 to Robin W. Baird, 14809-03 to Douglas P. Nowacek, and 16239 to Daniel T. Engelhaupt. Photo identification was conducted under National Marine Fisheries Service general authorization letter of confirmation number 19903 to Andrew J. Read. All activities were approved by Institutional Animal Care and Use Committees at the respective institutions.

## Additional Files

Additional file 1 — Supplementary_Table_1_settings.csv

Detailed settings parameters for SPLASH10 tags.

